# The Changing Immune Landscape of Innate-like T Cells and Innate Cells Throughout Life

**DOI:** 10.1101/2024.08.14.608020

**Authors:** Marziyeh Taheri, Christopher Menne, Jeremy Anderson, Shuo Li, Stuart P Berzins, Paul V Licciardi, Thomas M Ashhurst, Sedigheh Jalali, Daniel G Pellicci

## Abstract

Spectral flow cytometry is an advanced immunological tool that enables comprehensive analysis of the immune system by simultaneously comparing innate and adaptive immune cells. Here, using a 40-colour antibody panel we advance our knowledge of innate and innate-like T cells by investigating chemokine receptors, activation and maturation markers not usually assessed on these populations and examine age-related effects to these immune cell subsets. We characterised phenotypic changes of peripheral blood mononuclear cells (PBMC) in three age groups: newborn (cord blood), adults aged 20-30 years, and adults aged 70-80 years, focusing on innate-like T cells and innate cells, including MAIT cells, NKT cells, γδ T cells, ILCs, and Natural Killer (NK) cells. We identify subsets of double-negative (DN) T cells (CD4^-^ CD8^-^) and CD161^+^ T cells that increase in an age-related manner and exhibit a phenotype similar to innate-like T cells, MAIT cells and γδ T cells. Innate-like T cell subsets express similar patterns of the chemokine receptors and maturation markers CCR4, CCR6, CD27, CD38, CD57 and CD45RA, and resemble memory subsets of conventional CD4^+^ T cells and CD8^+^ T cells. We could detect ILCs in all age ranges, although the frequency of ILC1, ILC2, and ILC3 subsets decreased with age. Notably, we identify the NK maturation marker, CD57, as a universal marker that defines ageing populations of innate and adaptive immune cells. This study enhances our understanding of the ontogeny of human immune cells, highlighting significant age-related changes in the frequency and phenotype of immune cells.

## INTRODUCTION

The immune system is made up of a network of cells that display substantial diversity in their frequency, phenotype and function. Disruptions in certain immune compartments are not only caused by human diseases but also by aging, so providing a detailed description of the healthy immune system during aging, can help identify changes caused by disease throughout life. Broadly, the immune system is subdivided into two arms, innate and adaptive immunity, and it is vital to understand the key lineages of each arm. This includes in-depth phenotypic analysis of naive versus memory, regulatory, and activation markers through the expression of surface molecules such as CD45RA, CD27, CD57, CD38, and chemokine receptors (i.e. CCR4, CCR6, CCR7, CXCR3, and CXCR5) in different tissues such as peripheral blood (1, 2). Recent advancements in multi-omics methodologies and flow cytometry have facilitated the simultaneous analysis of an enormous number of different immune cell populations and provide an unprecedented depth to the exploration of novel combinations of markers and cell subsets, e.g. the analysis of innate markers (NKG2A, CD56, CD57, CD161) on T cells.

Factors like genetics, environmental conditions, infection, and aging can have profound impacts on the functioning of the immune system (3–5). Research on age-related alterations of the immune system has been ongoing for many decades with a focus on T cell subsets and T cell responses (6–9). Utilizing cord blood and peripheral blood is a powerful approach to compare the immune system of newborns with that of adult people and provides valuable insight into the effects of aging on the immune system development in the blood. As individuals age, their immune system undergoes immunosenescence rendering older individuals more susceptible to various diseases, particularly cancer and infection (10, 11). In contrast, the immune cells from cord blood are thought to exhibit an immature phenotype and have a reduced capacity to proliferate, secrete fewer cytokines and have reduced T cell responses (12). Immune profiling of cord blood cells is vital as cord blood is a valuable reservoir of hematopoietic stem cells and can be instrumental for various haematological and non-haematological malignancies and disorders (13, 14).

Here, we employed high-dimensional spectral flow cytometry to profile the blood immune cells in three healthy age groups: (i) newborns (cord blood), (ii) 20-30 years old, and (iii) 70-81 years old. In a previous study, we showed profound changes to the composition of lymphocytes of peripheral blood from infancy to adulthood (15). While our focus was on the developmental trajectory beyond infancy, particularly childhood and schooling age, this study provides detailed investigations in subsets of innate and innate-like T cells, taking advantage of a 40-colour antibody panel that enables more in-depth analysis of these cell types. The use of cord blood permits analysis of the immune system that is devoid from environmental influences like microbial or vaccine exposure and thus represents a truly naive immune system. Furthermore, we now include markers that allow the identification of Innate-Lymphoid Cells (ILC)-1, -2, and -3. ILCs lack a T cell receptor (TCR) to recognise specific antigens and their subsets are defined by differential expression of CD117 (c-kit) and CD294 (CRTH2). ILCs have recently been described in human blood and other tissues (16–18), although few studies have examined ILCs in cord blood (17, 19, 20). They share properties with innate-like T cells, including the expression of transcription factors and specific cytokines (21–23). Innate-like T cells represent up to 10% of T cells in human blood, play important roles in the protection against microbial infections and include NKT cells, Mucosal-Associated Invariant T (MAIT) cells and gamma delta (γδ) T cells (24, 25). Less is known whether other populations of innate-like T cells exist and their contribution to human immunity, although our comprehensive analysis identifies T cell subsets that bare striking similarity to innate-like T cells that also change in frequency and phenotype throughout life.

## RESULTS

In order to investigate age-related changes to innate and innate-like T cells, we surveyed the healthy immune system in cord blood mononuclear cells (CBMC) and adult PBMCs from participants that were clustered into two age groups, younger adults aged 20-30 years, and older adults aged 70-81 years (Suppl. table 1). We employed multi-colour full spectrum flow cytometry using a 40 fluorochromes antibody panel modified from our recent studies (15, 26) that facilitates the simultaneous identification and phenotyping of several immune cell lineages. In addition to lineage markers to decipher the majority of T cell-, B cell-, NK cell-, ILC-, unconventional T cell-, monocyte-, and dendritic cell-subsets that are commonly found in CBMC and PBMC, it features several chemokine receptors as well as markers of memory formation and maturation (Suppl. table 2-3). To accommodate for the high complexity of the data generated and the plethora of distinct immune cell subsets, we performed unsupervised high-dimensional integration and analysis workflow using the Specter toolkit in R, presenting our data using Uniform Manifold Approximation and Projection (UMAP) (Fig. 1). There were clear differences in the immune cell composition between newborn CBMC as well as young and old adult PBMC. The most striking differences were the near absence of CD4 and CD8 memory T cells from cord blood as well as the decrease of naïve T cells during aging. Moreover, innate-like T cells subsets, Vδ2+ T and MAIT cells were drastically increased in the two adult age groups (Fig. 1). Overall, this defines the baseline immune status of a naïve immune system in newborns, as wells as in healthy adults and facilitates identification of age-related changes in immune cell composition. Based on the immune cell subsets identified by the unsupervised clustering, we then further characterised the frequency of innate and innate-like T cell subsets using chemokine and maturation markers expressed by innate and adaptive immune cells.

**Figure 1.**
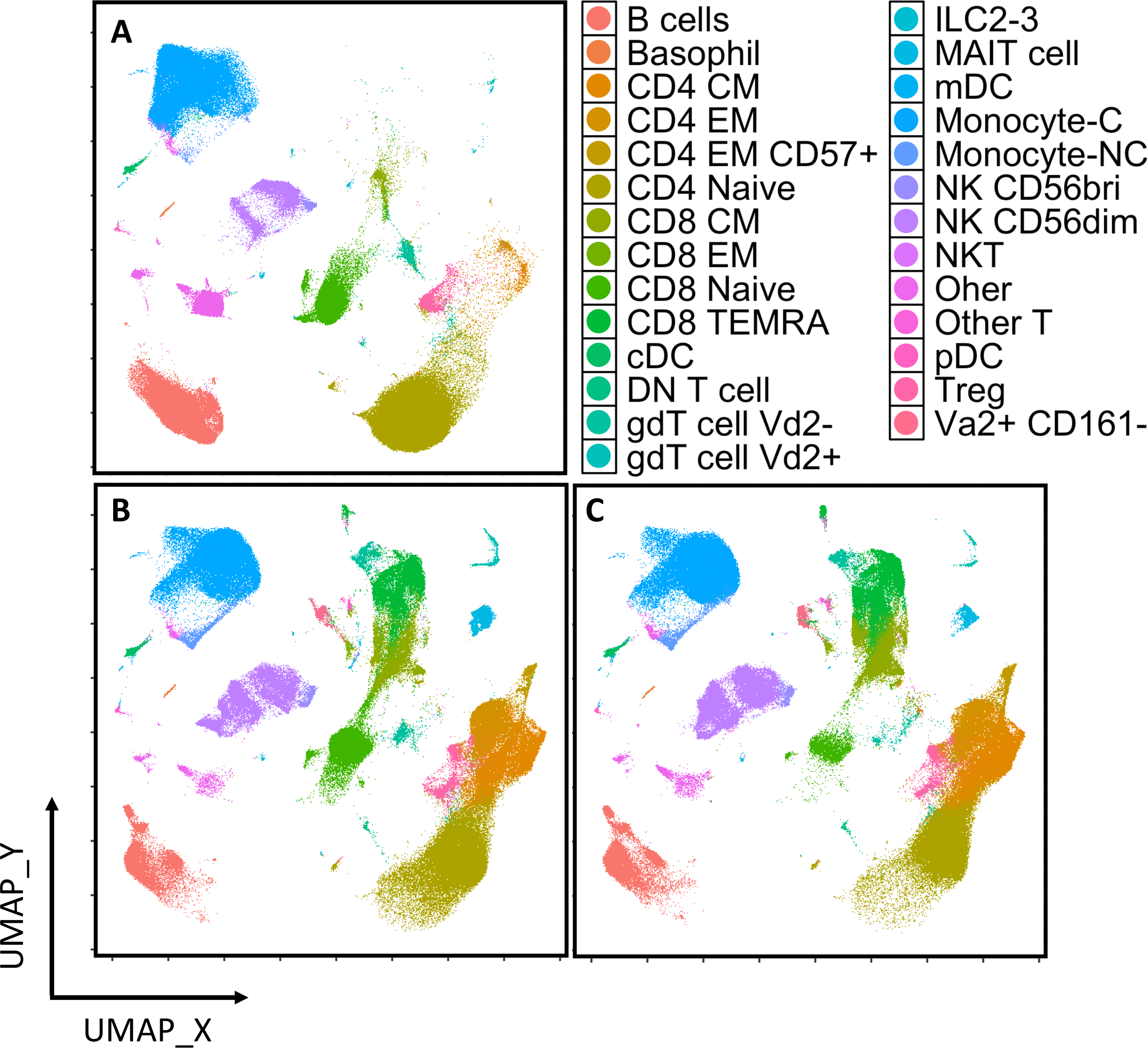
The transition of different immune cell subsets from peripheral blood mononuclear cells (PBMCs) of all participants in three age groups. (A) cord blood (B) adults aged 20-30 (C) adults aged 70-81 was validated with unsupervised dimension reduction using Uniform Manifold Approximation and Projection (UMAP).

### Identification of Innate Lymphoid Cells (ILCs) from cord and adult blood

First, we determined the frequency of ILC1, ILC2 and ILC3 subsets amongst CD3^-^CD19^-^ CD14^-^CD16^-^CD127^+^CD56^-/+^CD161^+^ cells (Fig. 2A). These ILC subsets were defined by their differential expression of c-kit and CRTH2 in the cord blood, young adult blood, and older adult blood (Fig. 2A, 2B). Our data showed that although the proportion of ILCs was low in all three age groups (<2% of CD3-cells), their frequencies were higher in cord blood compared to blood from adult groups (Fig. 2B). Given the frequency of ILC2 was low across all blood samples, it precluded further downstream analysis of these cells (Fig. 2B). Phenotypic comparison of ILC1 and ILC3 indicated that CCR6 expression was highest (∼10%) in young adults and declined in the older age group (∼5%) (Fig. 2C, 2D and Supplementary fig 1). CD27 was expressed by both ILC1 (15%) and ILC3 in cord blood (1%); however, an age-related decrease of CD27 was observed. The same age-related trend was observed for the expression of CD45RA for both ILC1 and ILC3 (from ∼80% to ∼65%) (Fig. 2C, 2D and Supplementary fig 1). CD38 expression on ILC3 increased with age, while ILC1 recorded a large increase in the maturation marker CD57 in older adults compared to cord blood (Fig. 2C, 2D and Supplementary fig 1).

**Figure 2.**
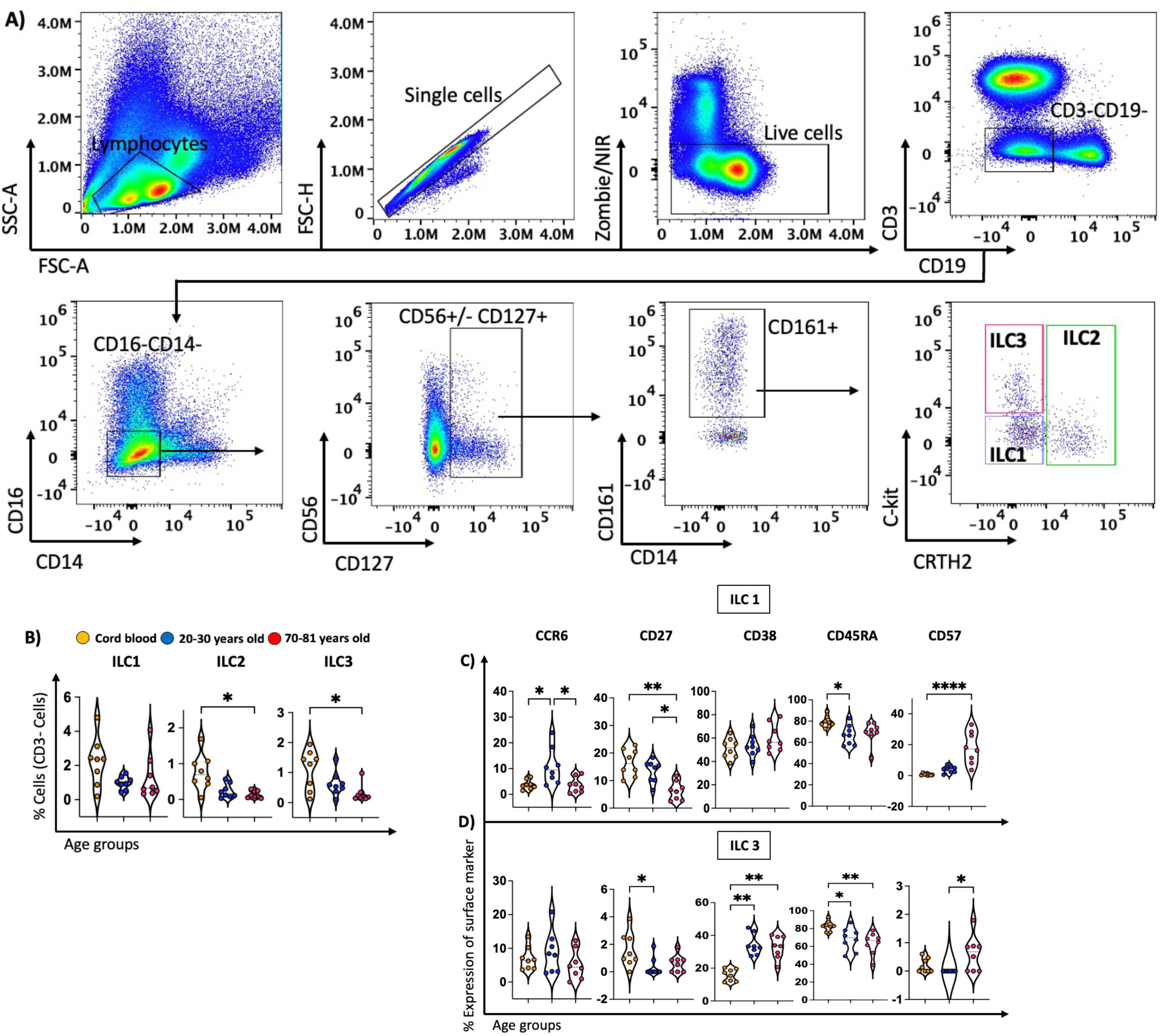
The proportion of different subsets of innate lymphoid cells (ILCs) decreased throughout life. (A) The flow cytometry plots show gating strategy for defining ILC subsets (ILC1, ILC2 and ILC3) from cord blood mononuclear cells (CBMCs) and peripheral blood mononuclear cells (PBMCs). Cells were gated for viable, single CD3-CD19-T cells and subsequently gated for CD16-CD14-CD56+/-CD127+CD161+ cells. Then ILC1, ILC2, and ILC3 subsets were defined using CD117(c-Kit) and CRTH2. (B) The violin plots show the age-related distribution of ILC1, ILC2, ILC3 within three age groups: cord blood (n=8, orange circles), young adult (n=8, 20-30 years old) (blue circles), and old adults (n=8, 70-81 years old) (red circles). (C, D). Violin plots representing the percentage of CCR6+, CD27+, CD38, CD45RA+ and CD57+ cells on ILC1 and ILC3 cells. Data are shown with the median. Each dot represents data from one individual donor and each colour represents one age group. A nonparametric Kruskal–Wallis test with Dunn’s multiple comparisons test was used for comparing all three groups. P-values are P > 0.05 (ns), *P ≤ 0.05; **P ≤ 0.01; ***P ≤ 0.001; ****P ≤ 0.0001.

### Characterisation of innate-like T cells

Analysis of MAIT cells and Vδ2^+^ γδ T cells showed minimal presence of these cells in cord blood. Their frequency increased in young adult blood (∼4%) but trended lower in older adults (∼1%) (Fig. 3A, 3B). Interestingly, the frequencies of NKT cells and Vδ2^-^ γδ T cells remained unchanged in cord blood and adult blood, irrespective of age (Fig. 3B).

**Figure 3.**
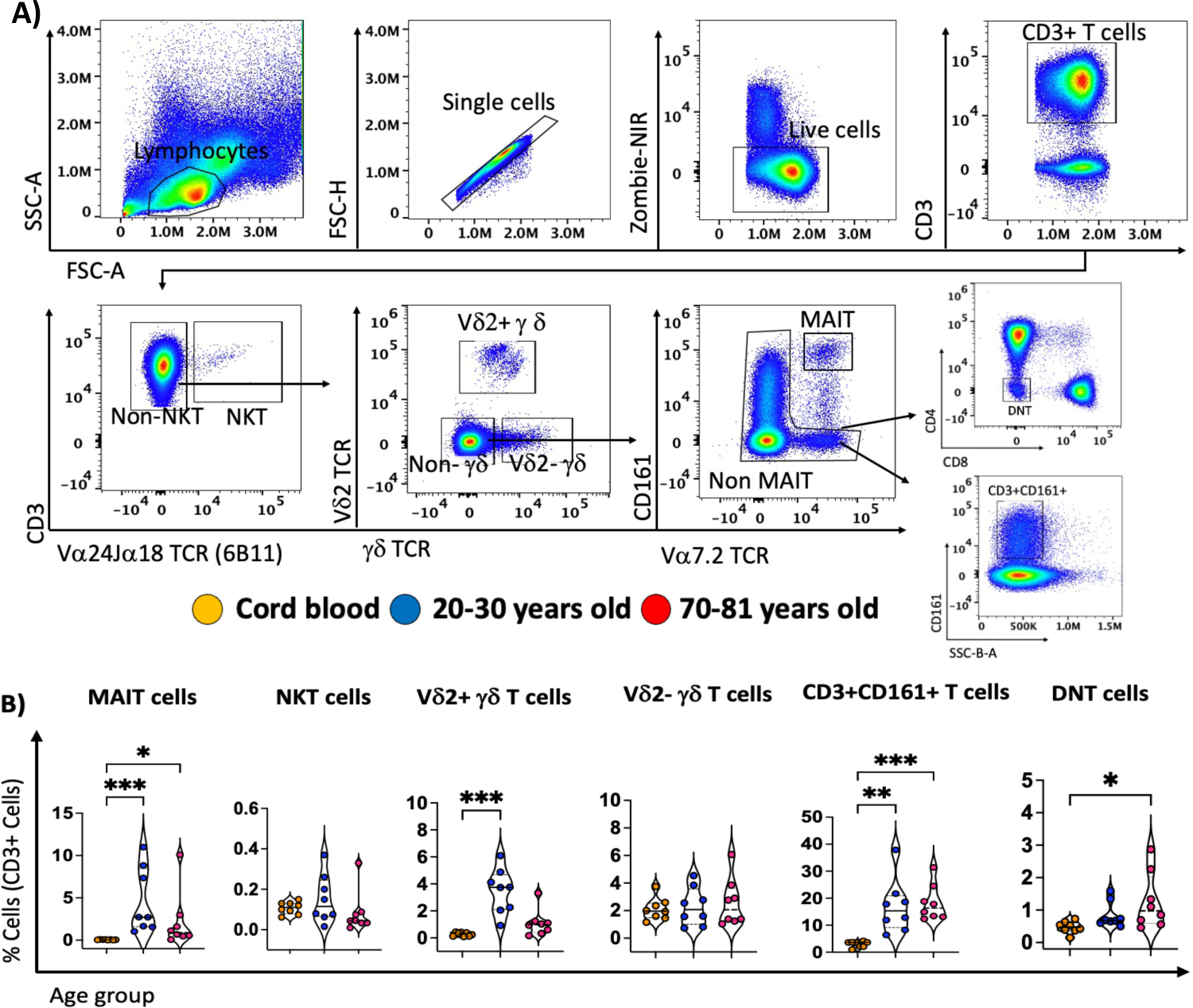
Variable alteration of the frequency of unconventional T cells by age. (A) The flow cytometry plots show gating strategy for defining different subsets of unconventional T cells including Mucosal-Associated Invariant T (MAIT) cells, Vδ2+ and Vδ2-gamma delta (γδ) T cells, Natural Killer T (NKT) cells, and two newly defined subsets: CD3+ CD161+ T cells and double negative (DN) T cells from cord blood mononuclear cells (CBMCs) and peripheral blood mononuclear cells (PBMCs). (B) The violin plots show the age-related distribution of unconventional T cells, CD3+ CD161+ T cells and double negative (DN) T cells within three age groups: cord blood (n=8, orange circles), young adult (n=8, 20-30 years old) (blue circles), and old adults (n=8, 70-81 years old) (red circles). Data are shown with the median. Each dot represents data from one individual donor and each colour represents one age group. A nonparametric Kruskal–Wallis test with Dunn’s multiple comparisons test was used for comparing all three groups. P-values are P > 0.05 (ns), *P ≤ 0.05; **P ≤ 0.01; ***P ≤ 0.001; ****P ≤ 0.0001.

After exclusion of these known subsets of innate-like T cells (NKT cells, MAIT cells and Vδ2^+^ γδ T cells), we observed large numbers of CD3+ T cells that still expressed CD161 (Fig. 3A, 3B), tentatively suggesting these cells represent another population of innate-like T cells. These CD3^+^CD161^+^ T cells expressed intermediate (int) levels of CD161 compared to MAIT cells and were virtually undetectable in cord blood but made up ∼15% of CD3^+^ T cells in adult blood (Fig. 3B). Similarly, after exclusion of known subsets of unconventional T cells, we also examined CD4^-^CD8^-^ T cells, herein defined as DN T cells (Fig. 3A) which revealed an age-related increased in their frequencies from cord blood to the adult age group (Fig. 3B).

To further understand the complexities of human innate-like T cell subsets, we compared the phenotype of MAIT cells and Vδ2^+^ γδ T cells with CD3^+^CD161^+^ T cells and DN T cells using a range of cell surface markers and chemokine receptors (Fig. 4 and Supplementary fig. 2-5). For example, CCR4, CCR7, and CD38 were highly expressed by MAIT cells and Vδ2^+^ γδ T cells and also by CD161^+^ T cells and DN T cells from cord blood (Fig. 4 and Supplementary fig. 2-5). CCR4 and CD38 showed an age-related decreasing trend on all of these T cell subsets (Fig. 4 and Supplementary fig. 2-5). The expression of CCR7 showed a similar age-related decrease, except for DN T cells on which it was highest in young adults (∼60%). Moreover, expression of CD45RA on these T cell subsets phenocopied CCR7 expression (Fig. 4 and Supplementary fig. 2-5). All four subsets of T cells expressed high levels of CD27 in cord blood (∼90%) and showed an age-related decrease which dropped to ∼40-50% in older adults (Fig. 4 and Supplementary figs. 2-5). Notably, CD57 expression increased with age, and it was the highest in DN T cells from older adults’ group (∼40%).

**Figure 4.**
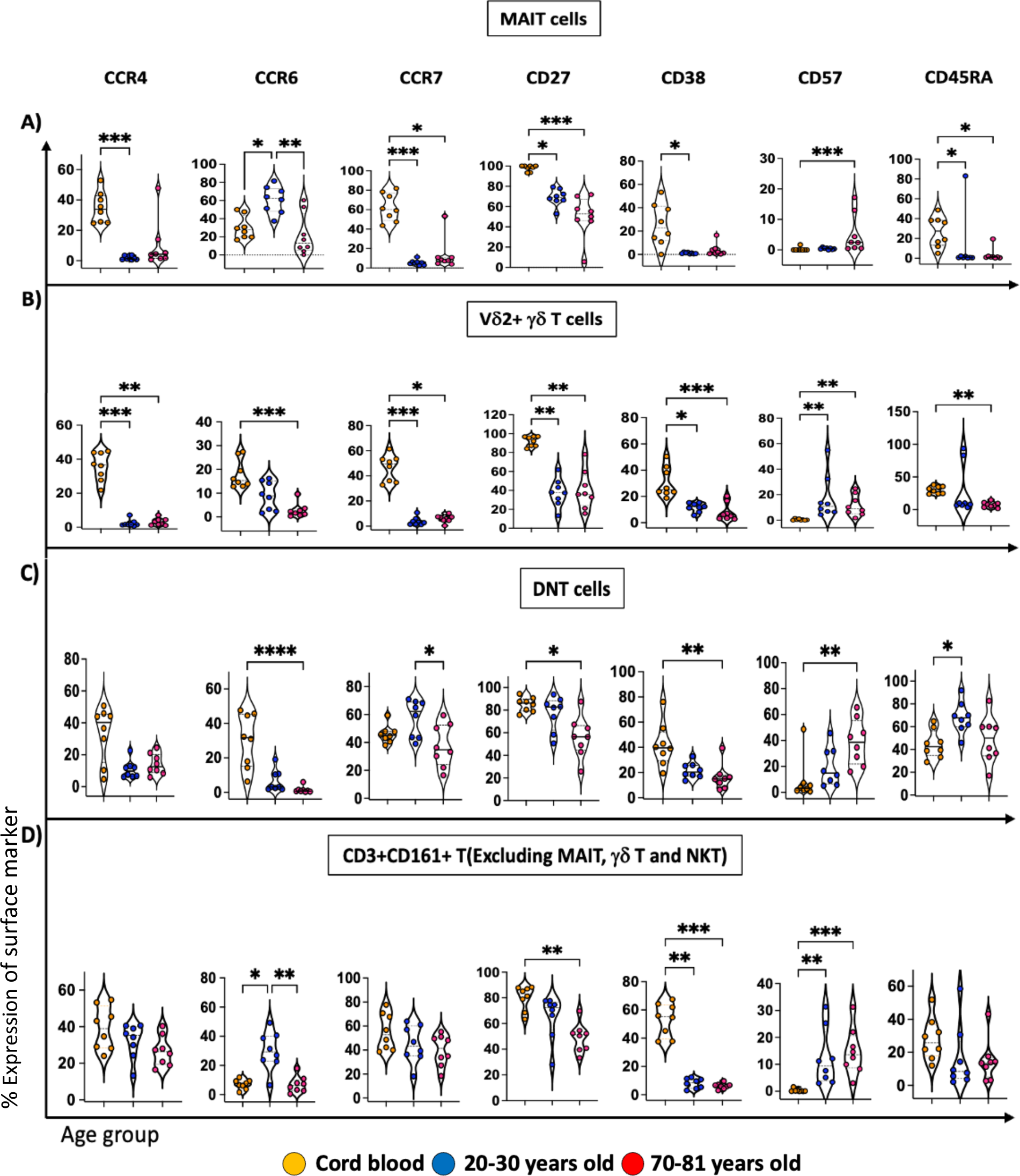
Comparison of cell surface marker expression indicate similarities between MAIT, Vδ2+γδ T, and two newly descried subsets of T cells: DNT and CD3+ CD161+ T cells. **(A-D)** The violin plots represent the age-related distribution of CCR4+, CCR6+, CCR7+, CD27+, CD38+, CD57+ and CD45RA+ cells on **(A)** MAIT cells, **(B)** Vδ2+γδ T, **(C)** DNT cells, and **(D)** CD3+CD161+ T cells. High dimensional flow cytometry was carried out on a total of 24 CBMCs/PBMC samples, cord blood (n=8, orange), young blood donors (n=8, blue) and older adult blood donors (n=8, red). Data are shown with the median. Each dot represents data from one individual donor and each colour represents one age group. A nonparametric Kruskal–Wallis test with Dunn’s multiple comparisons test was used for comparing all three groups. P-values are P > 0.05 (ns), *P ≤ 0.05; **P ≤ 0.01; ***P ≤ 0.001; ****P ≤ 0.0001.

CCR6 expression which is often used to define type III effector populations of Vδ2+ γδ T cells that express genes required for IL-17 production (27) trended similar between MAIT cells and CD3^+^CD161^+^ cells where it was low in cord blood (10-20%), increased in young adults (30-60%), and decreased in older adults (∼10%). The expression of this chemokine receptor was also similar between Vδ2^+^ γδ T cells and DN T cells where an age-related decrease from cord blood to adult was detected (from ∼20% to ∼2%) (Fig. 4 and Supplementary figs. 2-5).

While the frequency of Vδ2^-^ γδ T cells did not change with age, we observed the same phenotypic changes of this subset compared to other unconventional T cells in terms of an age-related decrease in the expression of CD27, CD38, CCR7, as well as CD28 and an increasing trend in the expression of CD57 in older individuals (Supplementary fig. 6). However, the expression of CCR4 trended opposite in Vδ2^-^ γδ T cells which increased over time, compared to the trend observed in other subsets of innate-like T cells (Fig. 4 and Supplementary figs. 2-5).

Taken together, our data highlights extraordinary age-related changes to the frequency and phenotype of innate-like T cells, and we identify similarities between these cells and CD3^+^CD161^+^ T cells and DN T cells.

### Unconventional T cell subsets share phenotypic properties with memory subsets of conventional T cells

A defining property of human innate-like T cells is that they exhibit maturation/memory markers that are typically acquired during development in the thymus (28–30). To determine if there were any similarities between the phenotype of innate-like T cells and conventional CD4+ and CD8+ memory T cell subsets, we compared the surface marker as well as chemokine receptor expression of MAIT cells and Vδ2^+^ γδ T cells to effector memory T cells (T_EM_), central memory T cells (T_CM_) and effector memory T cells expressing CD45RA (T_EMRA_) (Fig. 5A). Similar to our findings of the expression of CD27, CD38, and CD57 on innate like T cells (Fig. 4), we detected a decreasing trend in CD27, CD38 and an age-related increase in CD57 expression on all CD4^+^ and CD8^+^ memory T cell subsets (Fig. 5B). Moreover, CD38 was expressed at high levels on T_CM_ cells from cord blood (∼90%), while other T cell subsets (T_EM,_ T_EMRA,_ MAIT cells and Vδ2^+^ γδ T cells) from cord blood had moderate proportions of CD38^+^ cells (∼10-20%) (Fig. 4A, 4B, Fig5B, and Supplementary fig. 7-8). These findings represent the maturation status of both conventional and innate-like T cells throughout life.

**Figure 5.**
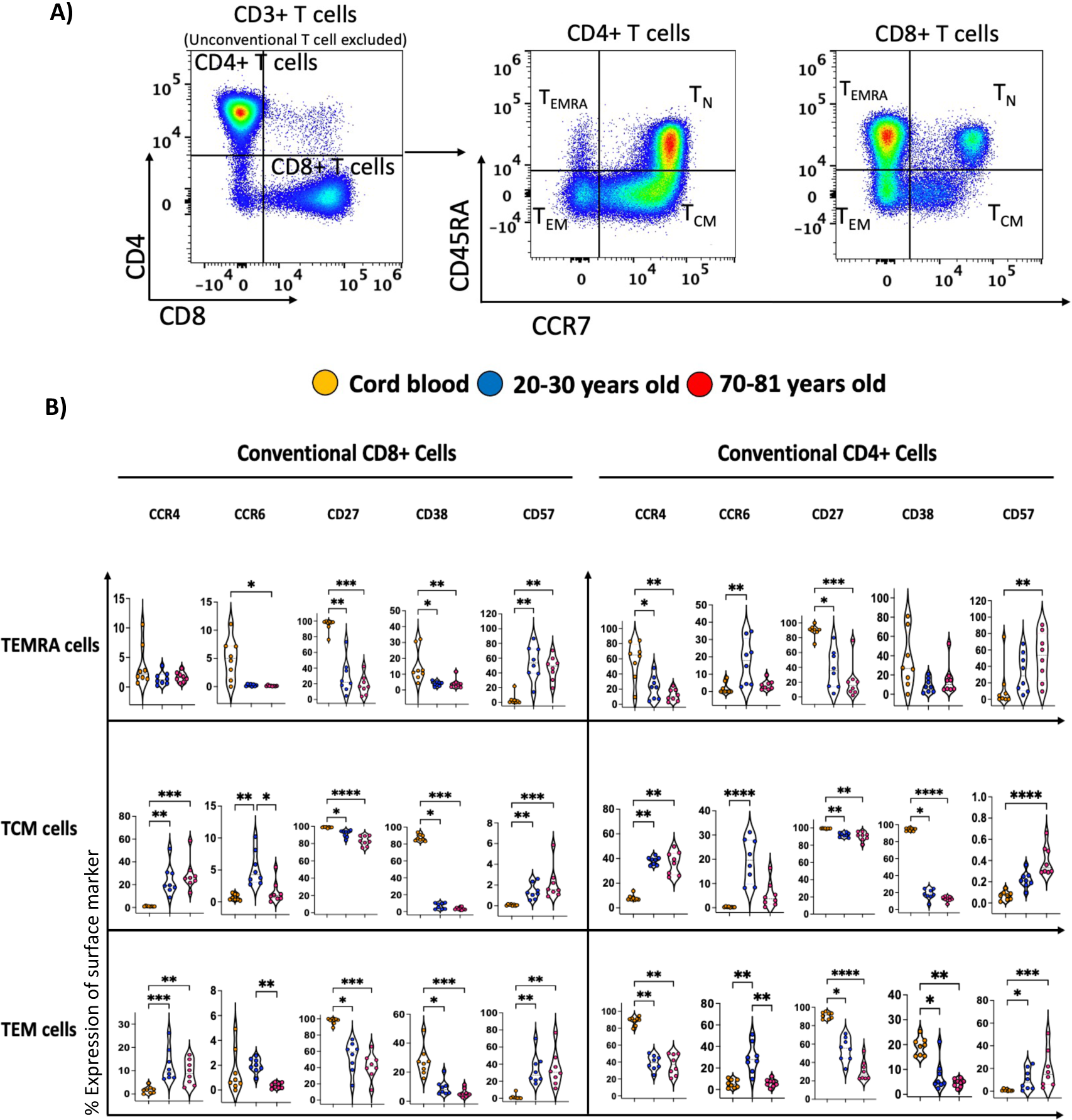
Memory subsets of conventional CD4+ T cells and CD8+ T cells exhibit some similar characteristics to unconventional T cells. **(A)** The flow cytometry plots show gating strategy for defining different memory subsets of CD4+ T cells and CD8+ T cells, after excluding NKT, γδ T cells, and MAIT cells from cord blood mononuclear cells (CBMCs) and peripheral blood mononuclear cells (PBMCs). **(B)** The Violin plots illustrated the age-related distribution of CCR4+, CCR6+, CD27+, CD38+, and CD57+ cells in T central memory (TCM), T effector memory (TEM), and T effector memory CD45RA+ (TEMRA) CD8+ T cells and CD4+ T cells within three age groups: cord blood (n=8, orange circles), young adult (n=8, 20-30 years old) (blue circles), and old adults (n=8, 70-81 years old) (red circles). The data are presented with the median, and each dot represents data from one individual donor. Each color represents a different age group. A nonparametric Kruskal–Wallis test with Dunn’s multiple comparisons test was used to compare all three groups. The p-values are as follows: P > 0.05 (ns), *P ≤ 0.05; **P ≤ 0.01; ***P ≤ 0.001; ****P ≤ 0.0001.

Like for MAIT cells and Vδ2^+^ γδ T cells from cord blood that had high frequencies of CCR4^+^ cells (∼40%) compared to adults’ blood (∼5%) (Fig 4), a similar trend was observed in CD4^+^ T_EM_ (from 60% to 10%) and T_EMRRA_ cells (from 90% to 40%) (Fig. 5B). We observed an opposite trend of the expression of CCR4 on other conventional memory T cells (CD4^+^ T_CM_, CD8^+^ T_CM_, CD8^+^ T_EM_) (Fig. 5B). Notably, the expression of CCR6 showed an age-related decreasing trend in CD8^+^ T_EMRA_ cells, which was similar to Vδ2^+^ γδ T cells (Fig. 4A, 4B, Fig5B, and Supplementary fig. 7-8). The rest of the memory subsets trended similar to MAIT cells, where the highest expression of CCR6 was observed in young adult blood (Fig. 4A, 4B, Fig5B, and Supplementary fig. 7-8).

### Innate-like T cells and innate cells undergo similar age-related changes in their phenotype

Vδ2^+^ γδ T cells (innate-like T cells) and NK cells (innate cells) share similar phenotypes and functions including the secretion of T_H1_ cytokines and cytotoxic killing granules (28, 31–33) and are thought to have complementary roles in human immune responses to microbes and cancer (32, 34). We closely examined the frequencies and phenotypes of two subsets of NK cells, CD56^dim^ and CD56^bright^, (Fig. 6A, 6B) and compared them to Vδ2^+^ γδ T cells. The frequency of CD56^bright^ NK cells and CD56^dim^ NK cells was low in cord blood and increased over time (Fig. 6B). By comparing CD56^dim^ and CD56^bright^ NK cells with Vδ2^+^ γδ T cells (Fig. 6C, 6D), we found that CD56^dim^ cells resembled Vδ2^+^ γδ T cells via expression of CD27, CD38 CD57 as well as CCR6 and CCR7 (Fig. 6D). More specifically, these cells typically express higher levels of CCR6, CCR7, CD27, CD38 and lower levels of CD57 in cord blood compared to cells from adult blood samples (Fig. 6D and Supplementary fig. 9). However, we observed the opposite trend when analysing the NK inhibitory marker NKG2A, between these two subsets. Specifically, the highest proportion of NKG2A^+^CD56^dim^ NK cells was detected in cord blood (∼70%) and it decreased with age (∼40%), while the lowest proportion of NKG2A^+^Vδ2^+^ γδ T cells was detected in cord blood (∼2%) and increased with age (∼60%) (Fig. 6D and Supplementary fig. 10).

Analysis of CD56^bright^ NK cells showed this subset of innate cells had some overlap with Vδ2^+^ γδ T cells and expressed the highest levels of CXCR3 in young adults’ blood compared to other age groups. CD56^bright^ NK cells also exhibited an age-related increase in the expression of CD57 (Fig. 6C) similar to innate-like T cells (Fig. 4 and Fig. 5). While the expression of CD161 on Vδ2^+^ γδ T cells trended higher in young adults’ blood compared to other age groups, it showed a decreasing trend in CD56^bright^ NK cells by age (from ∼95% to 60%) (Fig. 6C).

**Figure 6.**
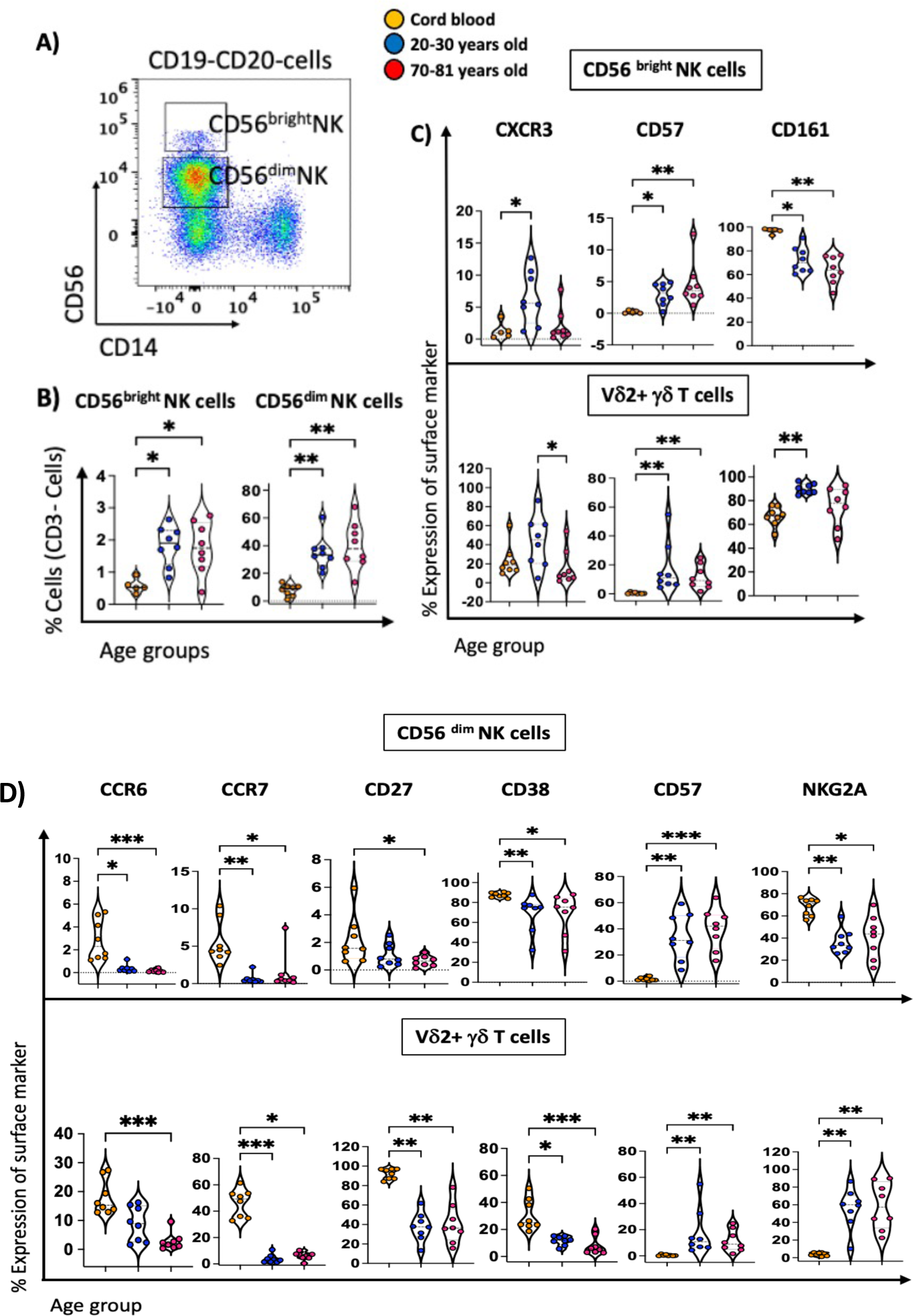
The phenotypic changes of Vd2+ gd T cells exhibit some similar characteristics to CD56^dim^ and CD56 ^bright^ NK cells. **(A)** Flow cytometric gating strategy for defining CD56^dim^ and CD56^bright^ NK cells from cord blood mononuclear cells (CBMCs) and peripheral blood mononuclear cells (PBMCs). **(B)** Violin plots illustrate the age-related distribution of CD56^dim^ NK and CD56^bright^ NK cells within three age groups: cord blood (n=8, orange circles), young adult (n=8, 20-30 years old) (blue circles), and old adults (n=8, 70-81 years old) (red circles). (**C-D)** Violin plots depicting the proportion of (**C)** CXCR3+, CD57+, and CD161+ cells within CD56^bright^ NK cells and Vd2+ gd T cells in the three age groups, and the proportion of (**D)** CCR6+, CCR7+, CD27+, CD38+, CD57+, and NKG2A in CD56^dim^ NK cells and Vd2+ gd T cells in the three age groups. Data are shown with the median. Each dot represents data from one individual donor and each colour represents one age group. A nonparametric Kruskal– Wallis test with Dunn’s multiple comparisons test was used for comparing all three groups. P-values are P > 0.05 (ns), *P ≤ 0.05; **P ≤ 0.01; ***P ≤ 0.001; ****P ≤ 0.0001.

## DISCUSSION

Spectral flow cytometry is continuing to advance at a rapid rate, allowing for superior analysis of the immune system in health and disease settings. Here, we used this technology to characterize the immune cells from cord blood, and peripheral blood of 20-30-year and 70-81-year-old healthy adults. In this study, we used a single 40-colour panel (26) to dive deeper into the effects of age on the complexities of the immune system in blood by comparing the newborns’ immune system (using cord blood) with young adults and elderly people, including the analysis of chemokine receptors, memory, activation and inhibitory markers. The current study is complementary to our previous publication (15), as we delve deeper into age-related changes to human MAIT cells, γδ T cells, ILC subsets as well as DN T cells and CD161^+^ T cells.

In line with a study by Darboe et al. that analysed circulatory ILCs in the Gambian population (ages 5-73 years) (20), we found no significant age-related changes in the frequency of ILC subsets between the young and old adult age group. However, higher frequencies of ILC1,2, and 3 subsets have been reported in cord blood compared to PBMCs of children and adults (16, 17, 35), and we observed the same trend, particularly for ILC2 and ILC3.

Changes in the frequency between the three subsets of ILCs showed that ILC1 and ILC3 subsets had reduced CD27 and CD45RA expression in older adult blood compared to cord blood, whereas CD57^+^ ILC1 and CD57^+^ ILC3 trended higher in older adults. These markers are well-established markers of T cell maturation and differentiation (36, 37) but are less commonly described in innate lymphoid cell subsets. Interestingly, we also observed the expression of CD38 did not change on the surface of ILC1 and increased on ILC3 in adults compared to cord blood. In contrast, the expression of CD38 showed an age-related decrease in all other conventional memory T cells, innate-like T cells, and innate cells we analysed. Given the multitude of functions described for CD38 ranging from cell activation to migration and cytokine release (38–40), this finding suggests a unique role of CD38 expression in regulating ILC3-specific immune function. Collectively our findings are suggestive of age-related maturation and differentiation of ILCs, which may impact their ability to influence immune outcomes, although further studies are needed to confirm this.

We and others have previously reported age-related changes to unconventional T cell lineages, including MAIT cells and Vδ2^+^ γδ T cells (15, 41, 42). For example, a negative correlation of MAIT cells with age has previously been reported in adult blood (41, 43–46), and we also showed that MAIT cells and Vδ2^+^ γδ T cells are more frequent in young adults compared to older adults and cord blood. After excluding known subsets of innate-like T cells (MAIT, NKT and γδ T cells), we focused on the remaining CD3^+^ T cells to identify DN T cells and CD161^int^ T cells. Many studies have analysed DN T cells from healthy human blood, however, they did not exclude known subsets of innate-like T cells (47–49). Excluding these cells from the analysis of DN T cells may enhance our understanding of what these cells are and better understand their unique characteristics and functions. For example, a study of individuals from Havana revealed a significant decrease in the frequency of blood DN T cells in older adults aged between 51 and 80 years (48), which could be attributable to a decrease in innate-like T cells, particularly Vδ2^+^ γδ T cells, which are predominantly DN T cells (28). By excluding innate-like T cells, we observed an opposite trend whereby the frequency of DN T cells actually increases with age. The phenotypic analysis of DN T cells reveal that the phenotype of these cells is strikingly similar to MAIT and Vδ2^+^ γδ T cells within each age group (CCR4, CCR6, CD27, CD38 and CD57), although CD45RA remained relatively high on DN T cells from adult blood compared to MAIT and Vδ2^+^ γδ T cells. These data suggest that DN T cells may contain a currently undefined subset of innate-like cells. Therefore, further investigations are required to define their TCR repertoire, antigen specificity, transcriptional profile, their function and ultimately their role in the human immune system.

Our data reveals that humans contain large populations of T cells that express intermediate (int) levels of CD161that are distinct from known subsets of innate-like T cells described above. Previous studies from the Klenerman group indicated that distinct subsets of CD161^+^ T cells share a conserved transcriptional signature with other unconventional T cell subsets and can be activated by IL-12 and IL-18 (50, 51). Furthermore, the same group revealed that CD8^+^ CD161^int^ T cells shared phenotypic and functional characteristics typical of unconventional T cells, including the expression of the transcription factors PLZF, Tbet, and Eomes; cytotoxic killing granules perforin and granzyme B; and cytokines IFNγ and IL-2 (52). Interestingly, these CD8^+^ CD161^int^ T cells included cells specific to viral peptides that are not usually recognized by innate-like T cells (52). In this study we found that the frequency of CD161^int^ T cells increased with age, similar to DN T cells. Our data also reveal a higher expression of CCR6 on the surface of CD161^+^ T cells in young adults compared to cord blood, which may suggest these cells have T_H_17 effector functions (15). Although these CD161^+^ cells share an overlapping phenotype with well-defined subsets of innate-like T cells and DN T cells, the trend of highest CCR6 expression in young adults resembles MAIT cells but is different from DN T cells and Vδ2+ γδ T cells that showed an age-related decrease.

Both γδ T cells and NK cells are heterogeneous populations of cells in pathological and physiological conditions. Previous studies have revealed similar phenotypic and functional characteristics between NK cells and Vδ2^+^ γδ T cells, emphasising how Vδ2^+^ γδ T cells can bridge the gaps between innate and adaptive immune responses. For example, both cell types can produce interferon-gamma (IFN-γ) and tumour necrosis factor-alpha (TNF-α) following activation or through antibody-dependent cellular cytotoxicity (ADCC) (53–55). They also exhibit dual effects in combating microbial and viral infections, cancerous cells or in graft-versus-host disease (GVHD) (53–55).

We found that Vδ2^+^ γδ T cells shared higher phenotypic similarities with CD56^dim^ NK cells compared to CD56^bright^ NK cells. It includes the expression of CCR6, CCR7, CD27 as well as CD38 in all three age groups. CD57, a maturation marker for NK cells (56–58), was found on the surface of Vδ2^+^ γδ T cells with the similar age-related increase observed for NK cell subsets. Higher expression levels of CD57 in CD16^+^CD56^dim^ NK cells have been reported on more mature cells with a higher cytotoxic activity and elevated responsiveness to CD16 mediated stimulation (58–60) which could be of interest given the expression of CD16 on human Vδ2+ gamma delta T cells (61). In contrast, higher expression of CD57 on T cells may represent a marker of immunosenescence in healthy individuals, reflecting a state of terminal differentiation and reduced proliferative capacity (62–64); however, in the context of viral infections (i.e. HIV), the presence of these cells might also indicate a virus-specific subset of CD8+ T cells that is involved in responding to the infection (64–66). Further assessments are needed to understand how the increased level of CD57 in Vδ2^+^ γδ T cells and other innate-like T cells influences their function. Furthermore, analysis of NKG2Ain cord blood showed an opposite trend as it was barely detectable on Vδ2^+^ γδ T cells whereas it was highly expressed on CD56dim NK. In contrast, both cell types express high levels of this inhibitory marker on their surface in adult blood.

Changes relating to the expression of chemokine receptors could have profound effects on cell migration and immune responses. Several studies have highlighted the important roles of cord blood-derived immune cells for therapeutic purposes such as cell therapies, paediatric transplantation, metabolic diseases, and brain injury (67–70). Moreover, the acquisition of CD57 on several subsets of innate-like cells and memory T cells was observed in older adults which could be an indication of altered function in the aged population. Further functional and transcriptional analysis will provide more insight into poorly characterised populations of innate cells, such as DN T cells and may lead to greater therapeutic opportunities to treat human disease.

Altogether, our study underscores the dynamic nature of the immune system across different life stages, from the immature immune phenotype observed in cord blood to the aged, immunosenescent profile in older adults. By utilising high-dimensional spectral flow cytometry, we were able to identify and characterise age-related phenotypic changes in innate-like T cells, as well as innate cells. These findings not only deepen our understanding of immune development during life but also highlight the potential for identifying novel immune cell subsets that may play important roles in human immunity. This research lays the foundation for further exploration into targeted therapies and personalised medicine, particularly in designing age-specific vaccine strategies to enhance immune protection in the most vulnerable populations—newborns and the elderly.

## MATERIALS AND METHODS

### Ethics Statement

Healthy adult blood was provided by the Australian Red Cross Lifeblood, agreement number 23-06VIC-01 and cord blood was provided by the Royal Children’s Hospital, with ethics approval from the Royal Children’s Hospital Melbourne Human Research Ethics Committee (HREC24131).

### Human blood samples

Peripheral blood mononuclear cells (PBMCs) from adult donors and cord blood mononuclear cells (CBMCs) from umbilical cord blood were isolated using Ficoll-Paque and preserved in freezing media consisting of 10% dimethyl sulfoxide (DMSO) and 90% fetal bovine serum (FBS). Cells were frozen at a rate of -1°C/minute in a -80°C freezer using a Cool Cell (Corning) and then transferred to the vapour phase of liquid nitrogen (-196°C) for long-term storage. A total of 24 samples were examined in this study, 8 CBMC samples and 16 adult PBMC samples (8 young adult and 8 older adult samples). For adult blood, each group contained an equal ratio of male and female individuals (Suppl. table 1).

### High-dimensional flow cytometry analysis

A 40-colour antibody panel was used to assess the frequency of immune cell subsets (Suppl. table 3) in cord blood, young adult blood, and older adult blood, comparing their frequency and phenotypic properties. PBMC or CBMC samples were thawed and stained with an established antibody cocktail in three stages: (i) staining of chemokine receptors with antibodies was performed at room temperature for 20 minutes followed by (ii) surface markers staining at 4°C for 20 minutes, and then (iii) Zombie NIR to identify any dead cells, with four additional surface markers at 4°C for another 20 minutes. Cells were washed between each step by removing the supernatant after 5 minutes of centrifugation at 400g and resuspending the cell pellets with PBS + 2% FBS buffer. Cells were analysed using a 5-laser Cytek Aurora.

### Statistical analysis

FlowJo software (10.9.0 version; BD, USA) was used to apply a manual gating method, identifying all critical cell subsets of our experiment according to the gating strategy illustrated in Figures 2, 3, 5 and 6 and Supplementary Figures 7 and 8. Cell subsets with 50 events or less in flow cytometry were excluded from further analysis. To compare all three groups including cord blood, young and old blood from healthy adults, the nonparametric Kruskal–Wallis test with Dunn’s multiple comparisons test was employed and a adjusted P-value <0.05 was regarded as significant. Data were plotted using GraphPad Prism version 10 software (GraphPad Software Inc, San Diego, CA, USA), and a P-value < 0.05 was regarded as significant. In addition, we analysed our findings with unsupervised algorithms using the Spectre R toolkit as we described previously (15, 71).

## Author Contributions

MT performed the research, carried out data analysis, figure preparation and drafted the initial version of the manuscript. TMA contributed to data preparation. CM, JA, SL, SPB, PVL and TMA contributed to project ideas and edited the manuscript. SJ and DGP conceived the study, edited the final manuscript and supervised MT.

## Acknowledgements

DGP was previously supported by the CSL Centenary Fellowship (2019–2023) and is now supported by the Sylvia & Charles Viertel Fellowship. JA is supported by an NHMRC Investigator grant (2026573). This research is supported by NHMRC grant (2030186).

## Data Availability Statement

Most data in this study are present within the published article. Additional data are available by request to the corresponding authors.

## Supplementary figure legends

**Supplementary Figure 1.**
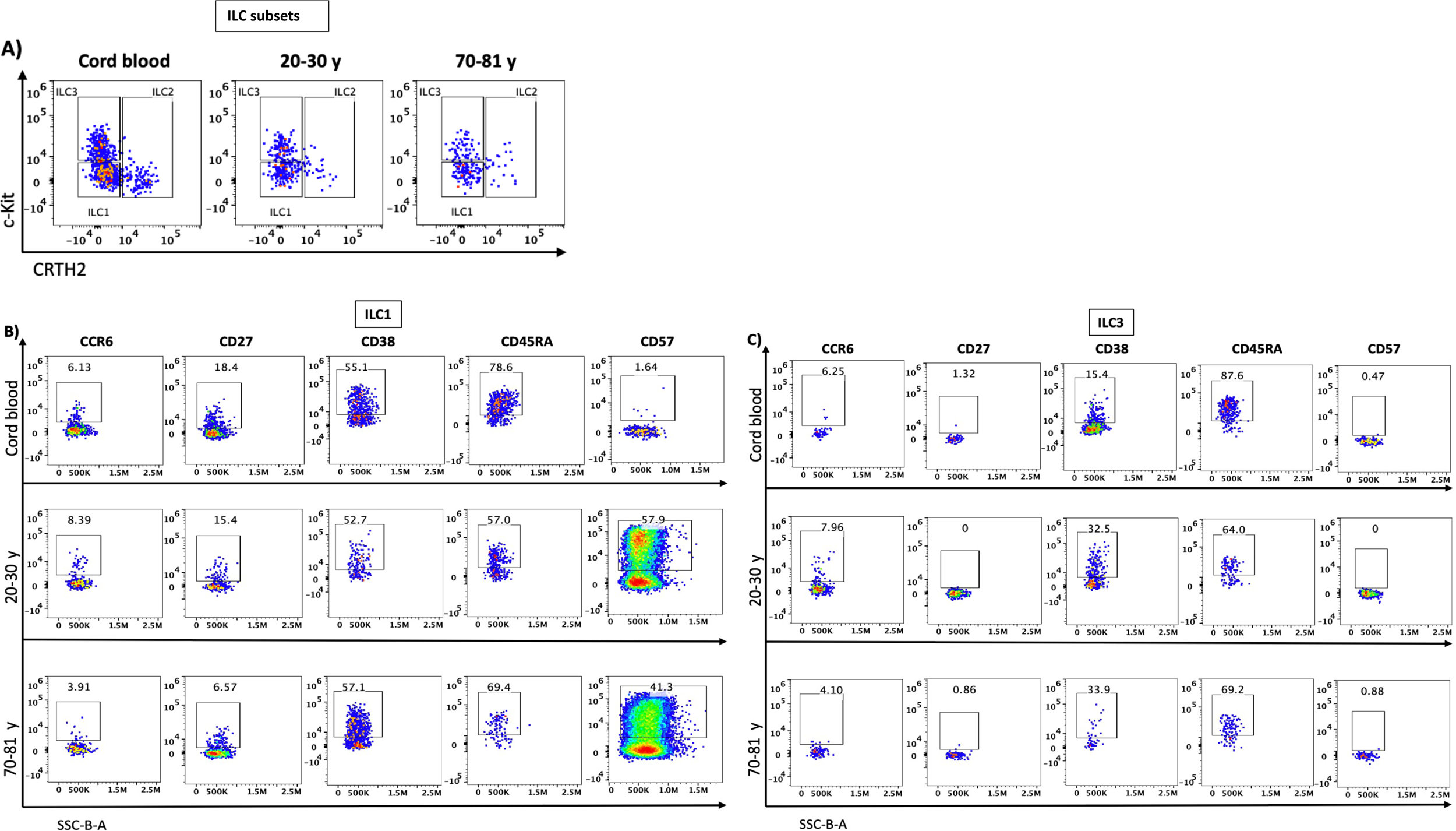
The phenotype of Innate Lymphoid Cells (ILCs) changed in adult blood compared to cord blood. (A) Flow cytometry dot plots depict the proportions of ILC1,2 and 3 with three age groups: cord blood, 20-30 years old, and 70-81 years old (B) The expression levels of CCR6, CD27, CD38, CD 45RA and CD57 on ILC1, (C) The expression levels of CCR6, CD27, CD38, CD45RA and CD57 on ILC1 and ILC3.

**Supplementary Figure 2.**
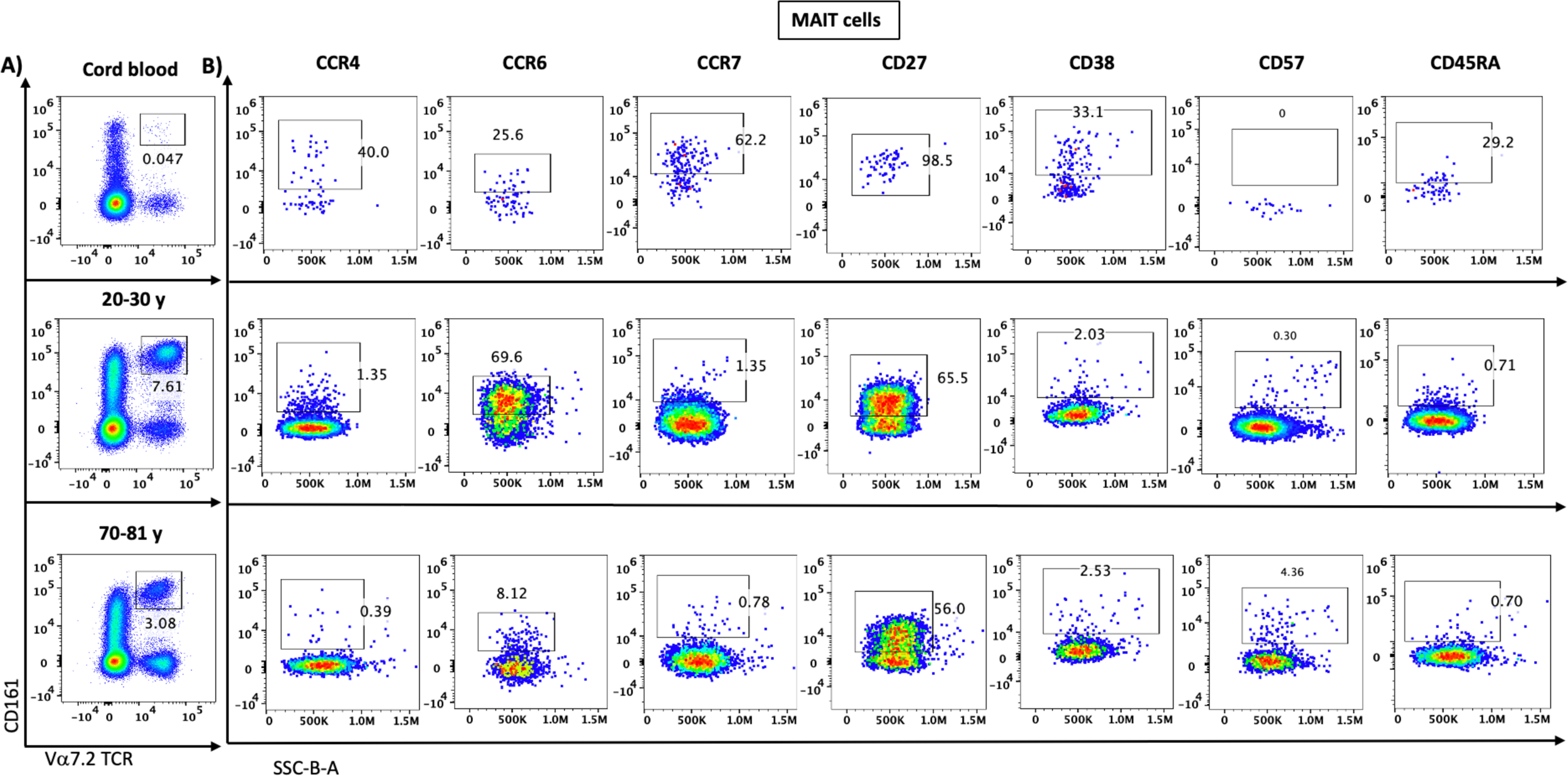
Changes in the phenotype of MAIT cells from cord blood to adult blood. **(A).** Flow cytometry dot plots depict the proportions of MAIT cells. **(B).** The expression levels of CCR4, CCR6, CCR7, CD27, CD38, CD57, and CD45RA on MAIT cells from cord blood, adult blood 20-30 years old age group, and adult blood 70-81 years old age group.

**Supplementary Figure 3..**
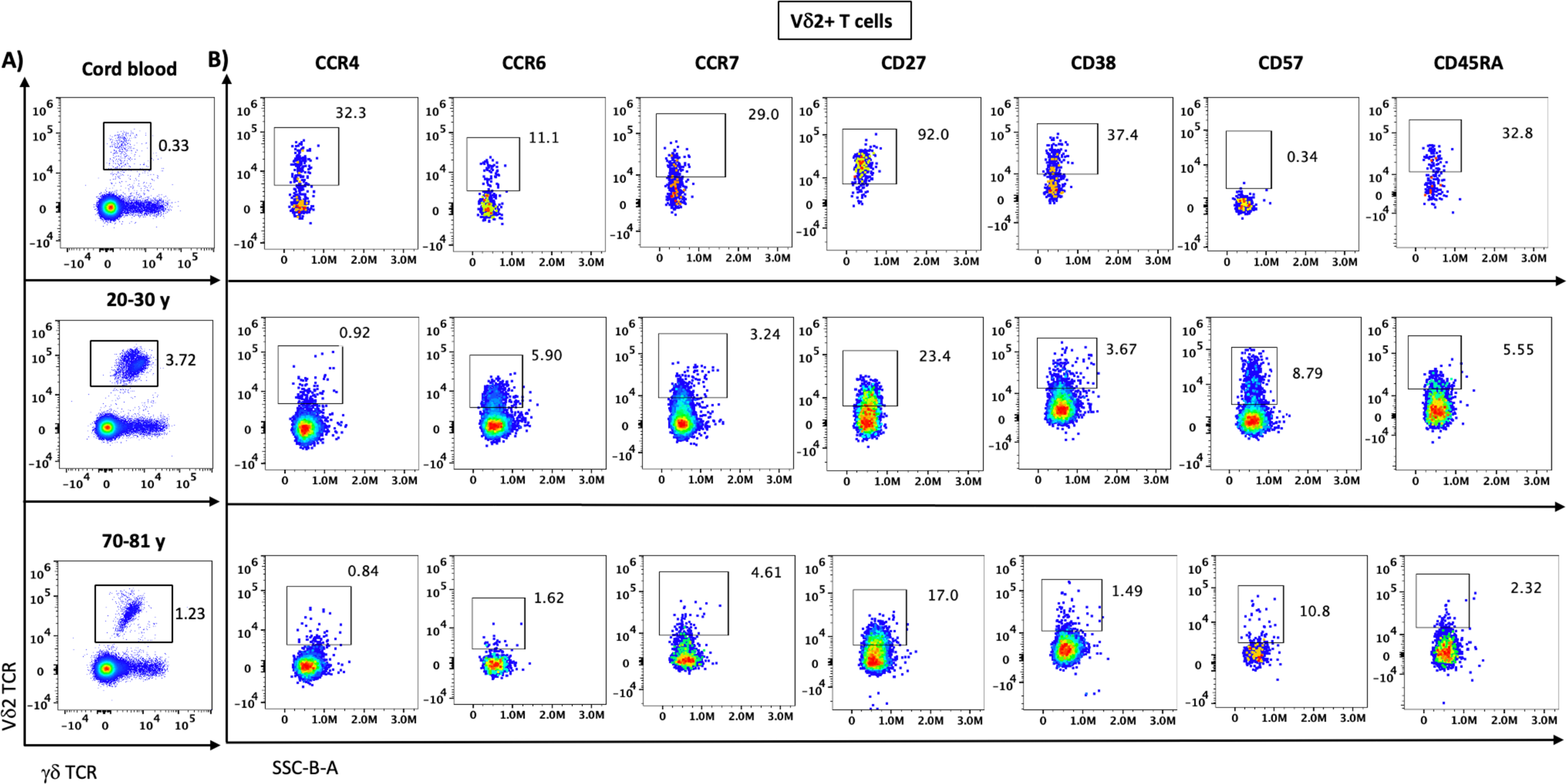
Changes in the phenotype of Vd2 gd T cells from cord blood to adult blood. **(A).** Flow cytometry dot plots depict the proportions of Vδ2+ γδ T cells. **(B).** The expression levels of CCR4, CCR6, CCR7, CD27, CD38, CD57, and CD45RA on Vδ2+ γδ T cells from cord blood, adult blood 20-30 years old age group, and adult blood 70-81 years old age group.

**Supplementary Figure 4.**
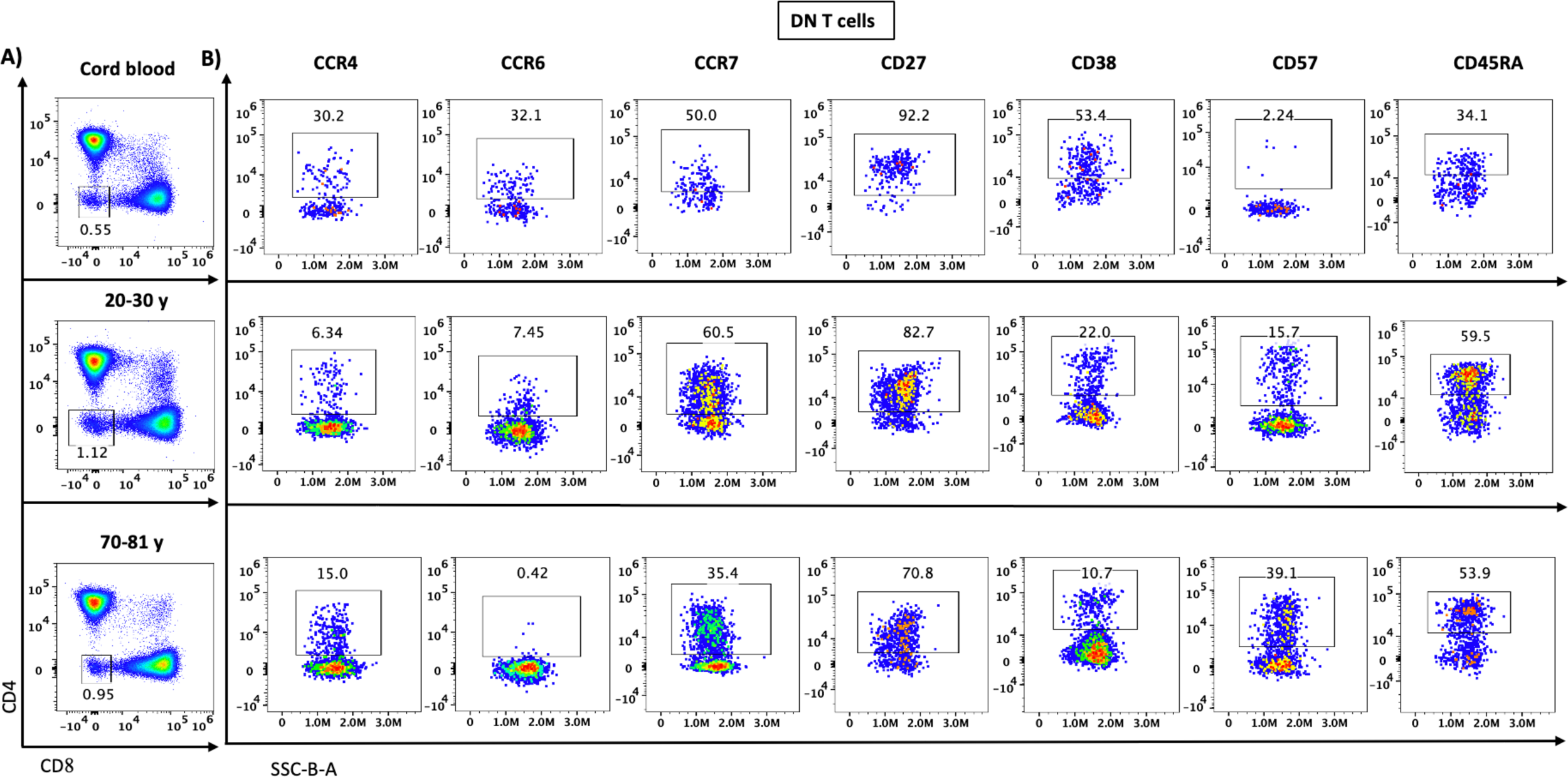
Changes in the phenotype of double negative (DN) T cells from cord blood and adult blood. **(A).** Flow cytometry dot plots depict the proportions of DN T cells, **(B).** The expression levels of CCR4, CCR6, CCR7, CD27, CD38, CD57, and CD45RA on DN T cells from cord blood, adult blood 20-30 years old age group, and adult blood 70-81 years old age group.

**Supplementary Figure 5.**
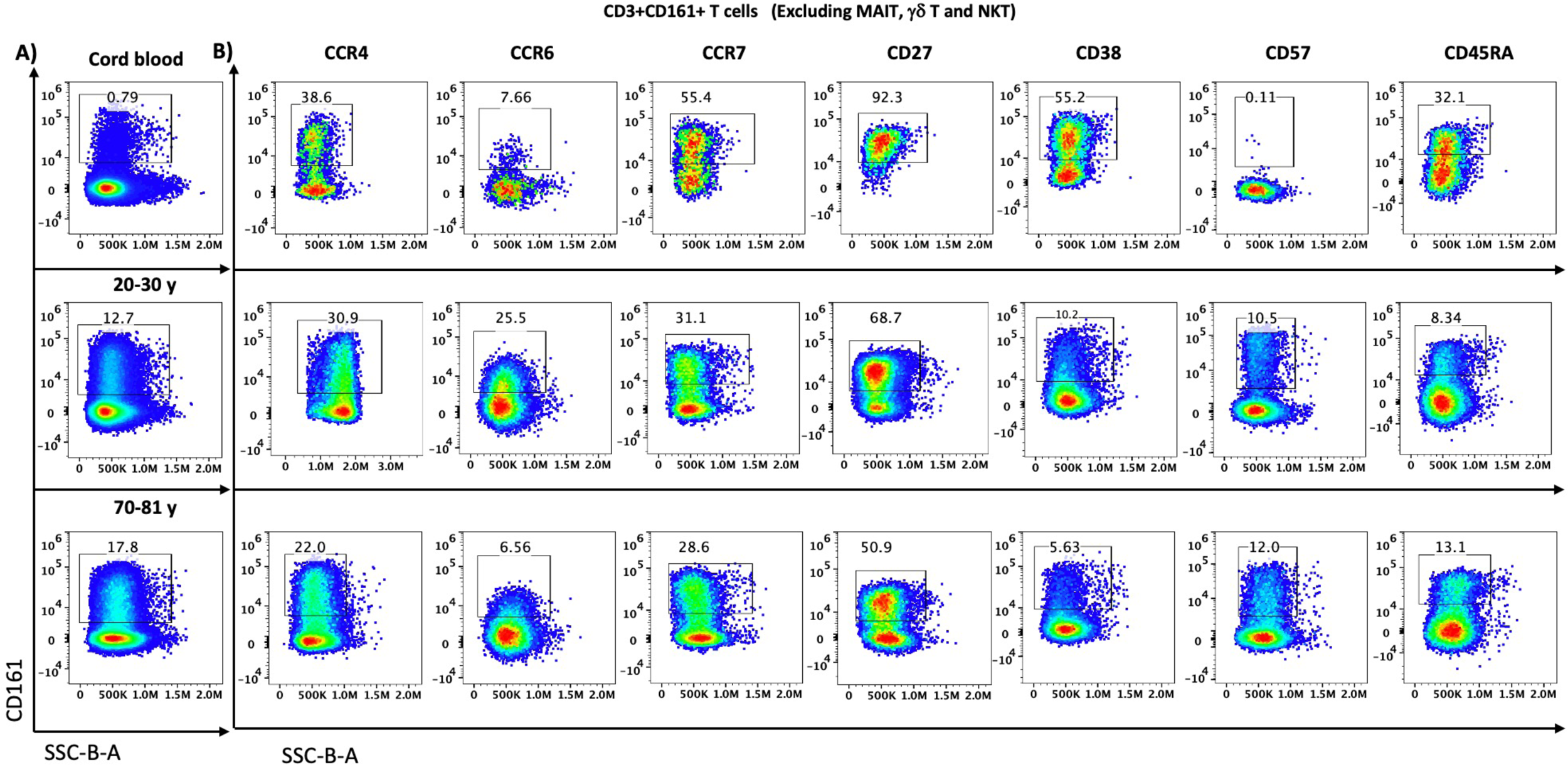
Changes in the phenotype of CD3+ CD161+ T cells from cord blood and adult blood. **(A).** Flow cytometry dot plots depict the proportions of CD3+CD161+ T cells. **(B).** The expression levels of CCR4, CCR6, CCR7, CD27, CD38, CD57, and CD45RA on CD3+CD161+ T cells from cord blood, adult blood 20-30 years old age group, and adult blood 70-81 years old age group.

**Supplementary Figure 6.**
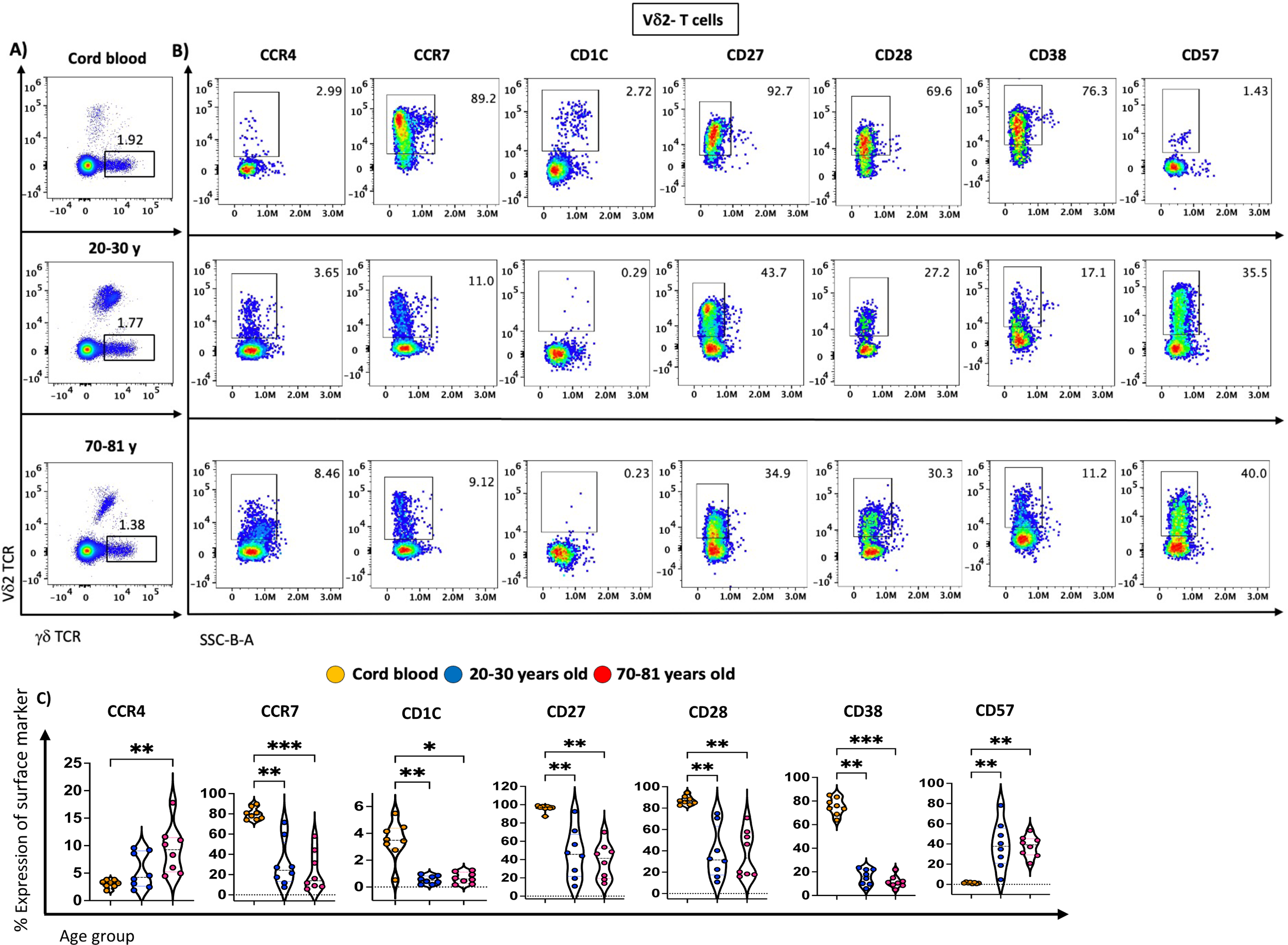
Changes in the phenotype of Vd2 - gd T cells from cord blood and adult blood. **(A).** Flow cytometry dot plots depict the proportions of Vδ2-γδ T cells **(B).** The expression levels of CCR4, CCR7, CD1C, CD27, CD28, CD38 and CD57 on Vδ2-γδ T cells from cord blood, adult blood 20-30 years old age group, and adult blood 70-81 years old age group. **(C)**. Violin plots show age-related distribution of surface markers including CCR4, CCR7, CD1C, CD27, CD28, CD38 and CD57 on Vδ2-γδ T cells. A total of 24 CBMC/PBMC samples, cord blood (n=8, orange), young adult blood (n=8, blue) and older adult blood (n=8, red). Data are shown with the median. Each dot represents data from one individual donor and each colour represents one age group. A nonparametric Kruskal–Wallis test with Dunn’s multiple comparisons test was used for comparing all three groups. P-values are P > 0.05 (ns), *P ≤ 0.05; **P ≤ 0.01; ***P ≤ 0.001; ****P ≤ 0.0001.

**Supplementary Figure 7.**
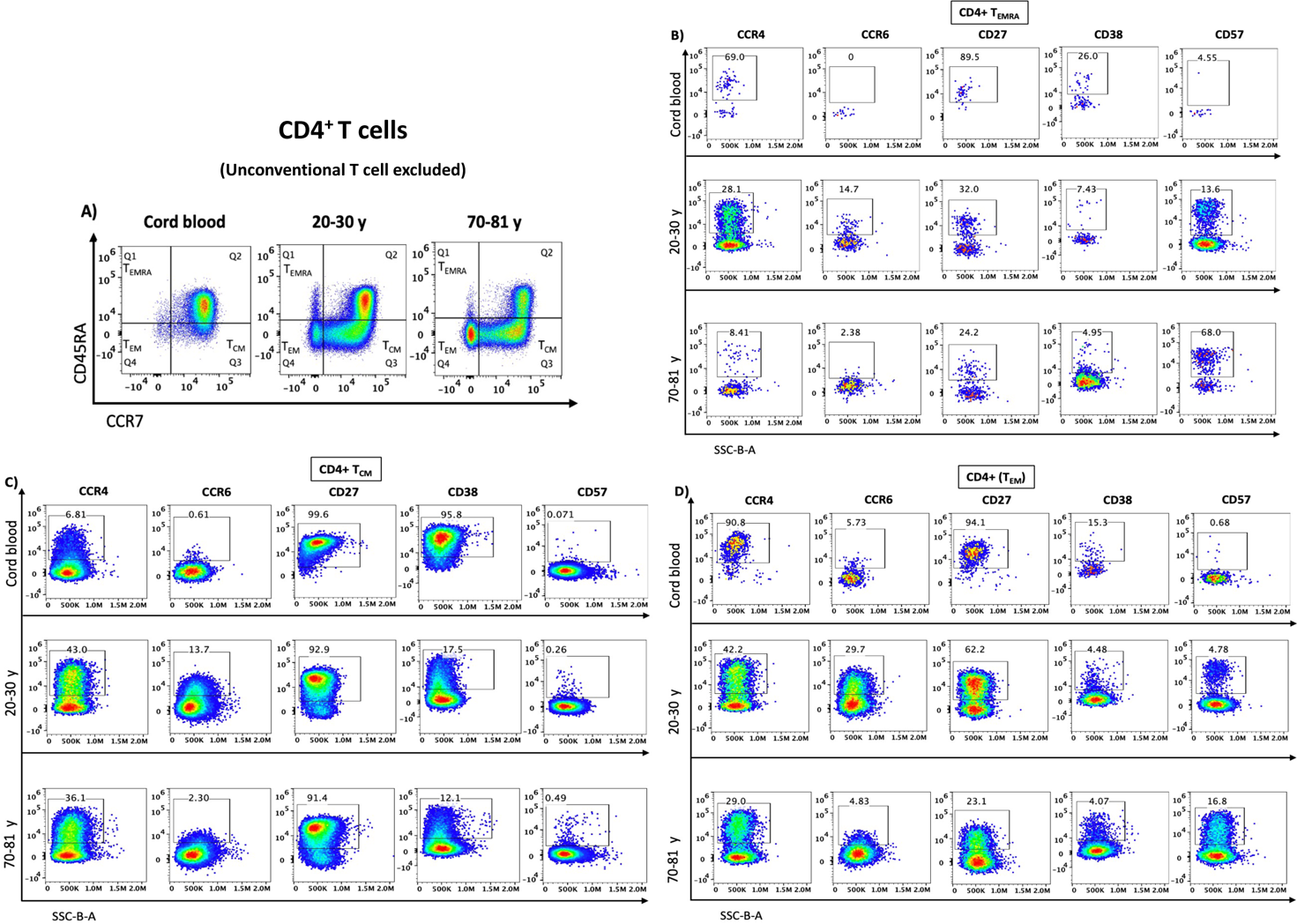
Changes in the phenotype of memory populations of CD4^+^ T cells from cord blood and adult blood. **(A).** Flow cytometry dot plots depict the proportions of T central memory (T_CM_), T effector memory (T_EM_) and T effector memory CD45RA +(T_EMRA_) CD4+ T cells. **(B-C).** The expression levels of CCR4, CCR6, CD27, CD38 and CD57 on **(B).** T effector memory CD45RA (T_EMRA_), **(C).** T central memory (T_CM_) and **(D)** T effector memory (T_EM_) CD4^+^ T cells from cord blood, adult blood 20-30 years old age group, and adult blood 70-81 years old age group.

**Supplementary Figure 8.**
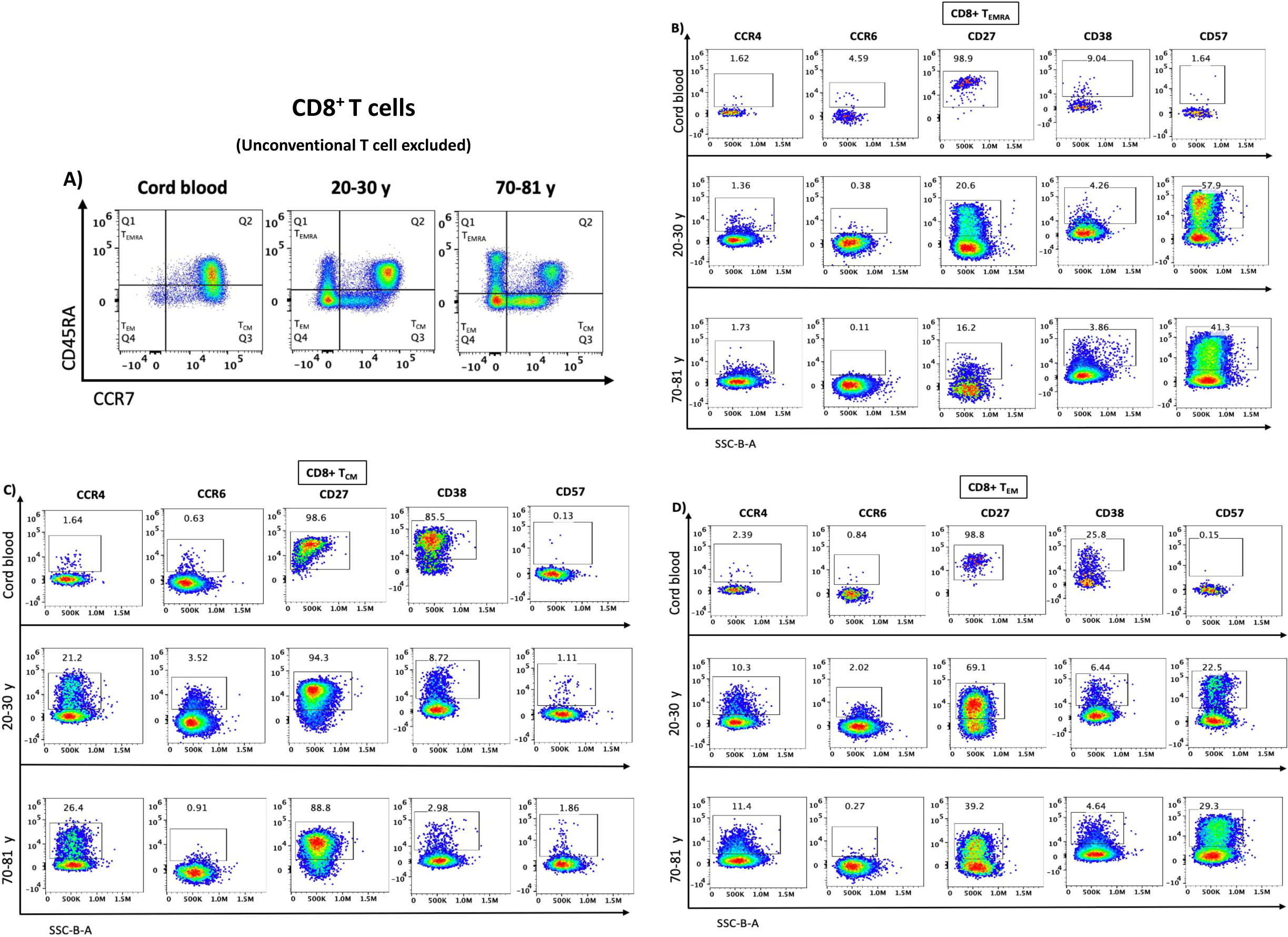
Changes in the phenotype of memory populations of C8^+^ T cells from cord blood and adult blood. **(A).** Flow cytometry dot plots depict the proportions of T central memory (T_CM_), T effector memory (T_EM_) and T effector memory CD45RA +(T_EMRA_) CD8^+^ T cells. **(B-C).** The expression levels of CCR4, CCR6, CD27, CD38 and CD57 on **(B).** T effector memory CD45RA (T_EMRA_), **(C).** T central memory (T_CM_) and **(D)** T effector memory (T_EM_) CD8^+^ T cells from cord blood, adult blood 20-30 years old age group, and adult blood 70-81 years old age group.

**Supplementary Figure 9.**
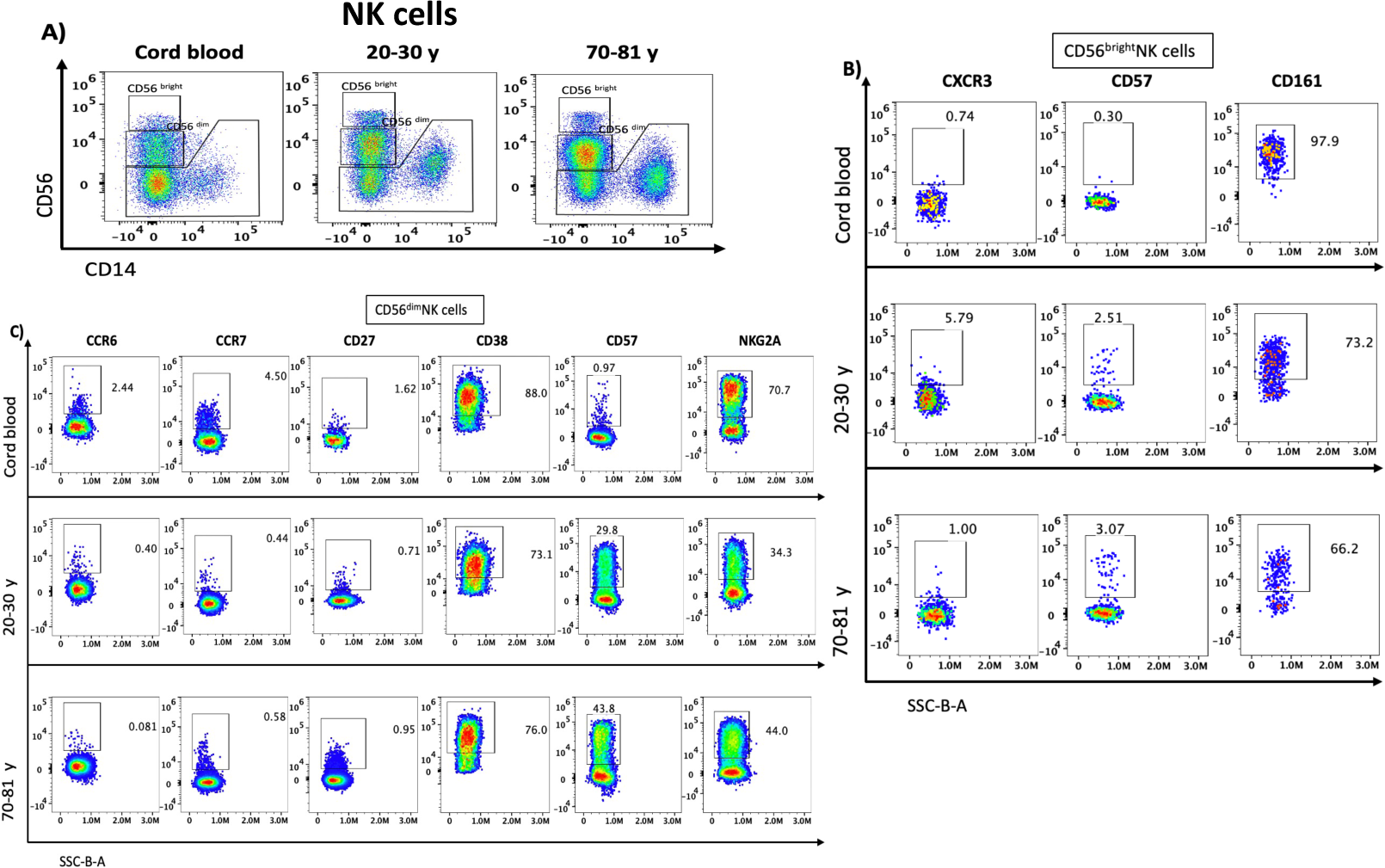
Changes in the phenotype of CD56 ^dim^ NK cells, and CD56 ^bright^ NK cells from cord blood and adult blood. **(A).** Flow cytometry dot plots depict the proportions of **CD56 ^dim^ NK cells, and CD56 ^bright^ NK cells (B).** and adult blood 70-81 years old age group.

**Supplementary Figure 10.**
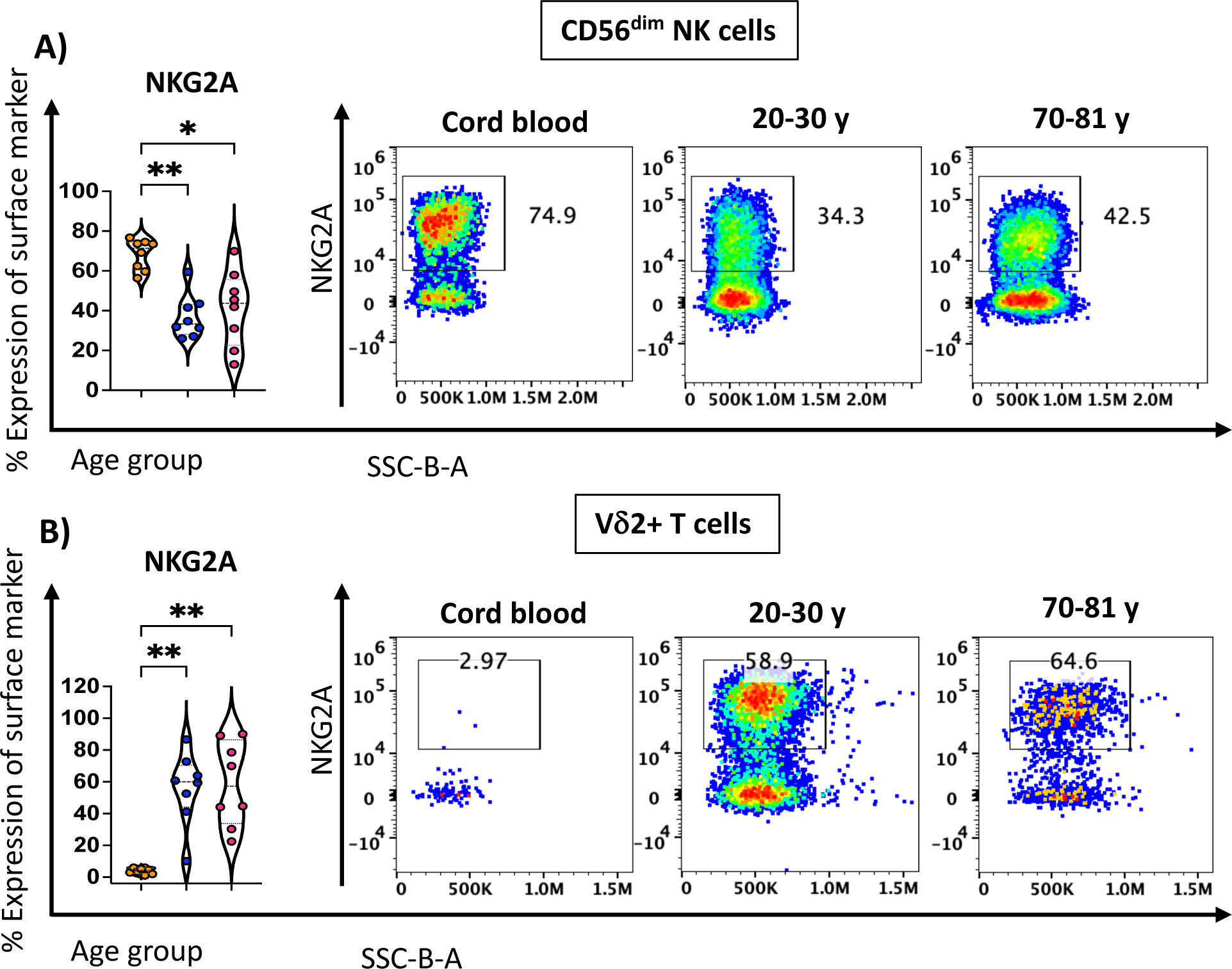
CD56^dim^ NK cells and Vd2+ T cells displayed a noticeably adverse expression level of NKG2A from cord blood to old adults. **(A).** Violin plots show the age-related distribution of NKG2A on CD56^dim^ NK cells and flow cytometry dot plots depict the expression levels of NKG2A on CD56^dim^ NK cells from cord blood, adult blood 20-30 years old age group, and adult blood 70-81 years old age group. **(B).** Violin plots show the age-related distribution of NKG2A on Vδ2+ T cells and flow cytometry dot plots depict the expression levels of NKG2A on Vδ2+ T cells from cord blood, adult blood 20-30 years old age group, and adult blood 70-81 years old age group.

---

**Supplementary Table 1:**

| Groups | Age range | No. of samples/Gender |
| --- | --- | --- |
| Group-1 | Cord blood | N: 8 |
| Group-2 | 20-30 Years | N: 8 (Male: 4/ Female: 4) |
| Group-3 | 70-81 Years | N: 8 (Male: 4/ Female: 4) |

**Supplementary Table 2:** Immune cell markers in cord blood and peripheral blood mononuclear cells (PBMCs).

| Cell populations | Markers |
| --- | --- |
| CD4<sup>+</sup> T cells | CD3+CD4+CD8-(Excluded MAIT, $\gamma\delta$ T, NKT) |
| CD8<sup>+</sup> T cells | CD3+CD8+CD4- (Excluded MAIT, $\gamma\delta$ T, NKT) |
| Double negative T cells | CD3+CD4-CD8-(Excluded MAIT, $\gamma\delta$ T, NKT) |
| CD3+CD161+ T cells | CD3+CD161+ (Excluded MAIT, $\gamma\delta$ T, NKT) |
| Naïve CD4<sup>+</sup> T cells | CD3+CD4+CCR7+CD45RA+ |
| CD4<sup>+</sup> T<sub>CM</sub> cells | CD3+CD4+CCR7+CD45RA- |
| CD4<sup>+</sup> T<sub>EM</sub> cells | CD3+CD4+CCR7-CD45RA- |
| CD4<sup>+</sup> T<sub>EMRA</sub> cells | CD3+CD4+CCR7-CD45RA+ |
| Naïve CD8<sup>+</sup> T cells | CD3+CD8+CCR7+CD45RA+ |
| CD8<sup>+</sup> T<sub>CM</sub> cells | CD3+CD8+CCR7+CD45RA- |
| CD8<sup>+</sup> T<sub>EM</sub> cells | CD3+CD8+CCR7-CD45RA- |
| CD8<sup>+</sup> T<sub>EMRA</sub> cells | CD3+CD8+CCR7-CD45RA+ |
| Vδ2<sup>+</sup> $\gamma\delta$ T cells | CD3+TCR$\gamma\delta$+Vδ2+ |
| Vδ2<sup>-</sup> $\gamma\delta$ T cells | CD3+TCR$\gamma\delta$+Vδ2- |
| MAIT cells | CD3+CD161+Vα7.2+ (Excluded $\gamma\delta$ T cells) |
| NKT cells | CD3+ Vα24JαQ TCR+ |
| CD56<sup>dim</sup> NK cells | CD3-CD19-CD20-CD14- CD56<sup>dim</sup> |
| CD56<sup>bright</sup> NK cells | CD3-CD19-CD20-CD14- CD56<sup>bright</sup> |
| Classical monocytes | CD3-CD19-CD20-CD56-CD14+CD16- |
| Intermediate monocytes | CD3-CD19-CD20-CD56-CD14+CD16+ |
| Non-classical monocytes | CD3-CD19-CD20-CD56-CD14-CD16+ |
| Dendritic cells (DCs) | CD3-CD19-CD20-CD56-CD14-CD16- |
| Myeloid DCs (mDCs) | CD3-CD19-CD20-CD56-CD14-CD16-CD11c+ |
| Plasmacytoid DCs (pDCs) | CD3-CD19-CD20-CD56-CD14-CD16-CD11c-<br>CD123+ |
| Innate lymphoid cells (ILC1) | CD3-CD19-HLA-DR-CD14-CD56-CD123-CD16-<br>CD161+CD127+CD25+CD117-CRTH2- |
| Innate lymphoid cells (ILC2) | CD3-CD19-HLA-DR-CD14-CD56-CD123-CD16-<br>CD161+CD127+CD25+CD117+CRTH2+ |
| Innate lymphoid cells (ILC3) | CD3-CD19-HLA-DR-CD14-CD56-CD123-CD16-<br>CD161+CD127+CD25+CD117+CRTH2- |

**Supplementary Table 3:** 40 colour Antibody panels for immune profiling of cord blood and peripheral blood mononuclear cells (PBMCs).

| **Immunogen** | **Conjugate** | **Manufacturer** | **Clone** |
| --- | --- | --- | --- |
| **Antibody stain 1** | **Room temperature** |  |  |
| **CD297 (PD-1)** | BUV615 | BD Biosciences | EH12.1 |
| **CD117 (c-Kit)** | AlexaFluor 647 | BioLegen | 104-D2 |
| **CD294 (CRTH2)** | BV510 | BD Biosciences | BM16 |
| **CD194 (CCR4)** | BV605 | BioLegend | L291H4 |
| **CD196 (CCR6)** | BV650 | BD Biosciences | 11A9 |
| **CD197 (CCR7)** | PE/Fire810 | BioLegend | G043H7 |
| **CD183 (CXCR3)** | APC | BD Biosciences | 1C6 |
| **CD185 (CXCR5)** | BUV805 | BD Biosciences | RF8B2 |
| **CD27** | PerCP-eFluor 710 | BD Biosciences | 0323 |
| **Antibody stain 2** | **4°C** |  |  |
| **CD3** | BUV395 | BD Biosciences | UCHT1 |
| **CD4** | BUV661 | BD Biosciences | SK3 |
| **CD28** | BUV563 | BD Biosciences | CD28.2 |
| **CD38** | BUV496 | BD Biosciences | HIT2 |
| **CD69** | BUV737 | BD Biosciences | FN50 |
| **CD56** | BV570 | BioLegend | HCD56 |
| **CD57** | Pacific Blue | BioLegend | HNK-1 |
| **CD141** | BV750 | BD Biosciences | 1A4 |
| **HLA-DR** | BV421 | BioLegend | L243 |
| **Vα7.2 TCR** | BV711 | BioLegend | 3C10 |
| **Vδ2** | BV480 | BD Biosciences | B6 |
| **iNKT 6B11** | BV785 | BioLegend | 6B11 |
| **γδTCR** | FITC | BD Biosciences | 11F2 |
| **IgM** | Spark blue 550 | BioLegend | MHM-88 |
| **CD11c** | PerCP | BioLegend | Bu15 |
| **CD45RA** | PerCP/cy5.5 | BD Biosciences | HI100 |
| **IgD** | PerCP/Fire806 | BioLegend | W18340F |
| **CD159a**<br>**(NKG2A)** | PE | Beckman Coulter | Z199 |
| **CD24** | PE-AF610 | BD Biosciences | SN3 |
| **CD1c** | PEcy5 | BioLegend | L161 |
| **CD16** | BYG710 | Cytek Biosciences | 3GB |
| **CD161** | PEvio770 | Miltenyi Biotec | 191 B8 |
| **CD8** | APC | BioLegend | SK1 |
| **CD19** | Spark NIR 685 | BioLegend | HIB19 |
| **CD127** | APC-R700 | BD Biosciences | HIL-7R-M21 |
| **IgG** | APC-H7 | BD Biosciences | G18-145 |
| **Antibody stain 3** | **4°C** |  |  |
| **CD14** | BB700 | BD Biosciences | M5E2 |
| **CD20** | Pacific orange | Invitrogen | HI47 |
| **CD25** | PE-CF594 | BD Biosciences | M-A251 |
| **CD123** | RY586 | BD Biosciences | 7G3 |
| **Zombie NIR** | NIR | BioLegend | Zombie NIR |

